# Assessing specificity testing in Lesion Network Mapping

**DOI:** 10.64898/2026.08.26.746668

**Authors:** Martijn P. van den Heuvel, Ilan Libedinsky, Sebastian Quiroz Monnens, Jonathan Repple, Luca Cocchi

## Abstract

Lesion Network Mapping (LNM) is a framework used for identifying symptom-related brain circuits by projecting lesion locations onto a normative connectome. Recent methodological investigations have raised concerns about the biological interpretation and specificity of the circuits derived using this method, with published LNM maps often showing high similarity across clinically unrelated conditions. *Specificity testing* has subsequently been put forward as the decisive step to ensure specificity to the symptom in question, accompanied by the argument that this step was not evaluated in the original methodological investigation. Yet, sensitivity testing, specificity testing, case-control LNM, permutation of group labels, and symptom-based LNM involve related operations on connectivity matrix *C*. We expand on specificity testing in LNM, clarify its relationship to other LNM steps and variants, and examine the persistent repetition among LNM specificity networks across studies. These considerations advance our understanding of the disease-specificity limitation of LNM and encourage the development of new methodological approaches for identifying brain circuits underlying psychiatric and neurological disorders.

## Introduction

Lesion Network Mapping (LNM) is a framework proposed to identify symptom and disease-associated brain networks by mapping lesion locations onto a normative functional connectome^1, 2^. Recent investigations^3, 4^ highlighted that LNM-derived networks are shaped by latent connectivity patterns, such that proposed networks may primarily reflect generic connectome structure rather than disease-specific circuitry. The ensuing discussion has focused on whether these concerns are resolved by *specificity testing*^5^, and more broadly on how disease specificity should be evaluated in LNM-derived networks.

### Repetition of LNM networks

An overview of 200 LNM studies^3^ highlighted substantial overlap among reported LNM networks, often with examples of r = 0.6–0.8 between unrelated conditions. This repetitiveness challenges the core aim of LNM to delineate symptom-specific brain networks. For example, a circuit reported to differentiate dysphoric-anxiosomatic symptoms in depression^6^ and recently proposed for treatment targeting of these symptoms^7^, closely resembles a network reported elsewhere for psychosis^8^, a DBS network associated with cognitive decline in patients with Parkinson’s disease^9^, as well as a network independently proposed to be associated with PTSD^10^ (Figure 1, |r| > 0.7; specificity LNM maps downloaded from the original publications without additional analysis).

**Figure 1.**
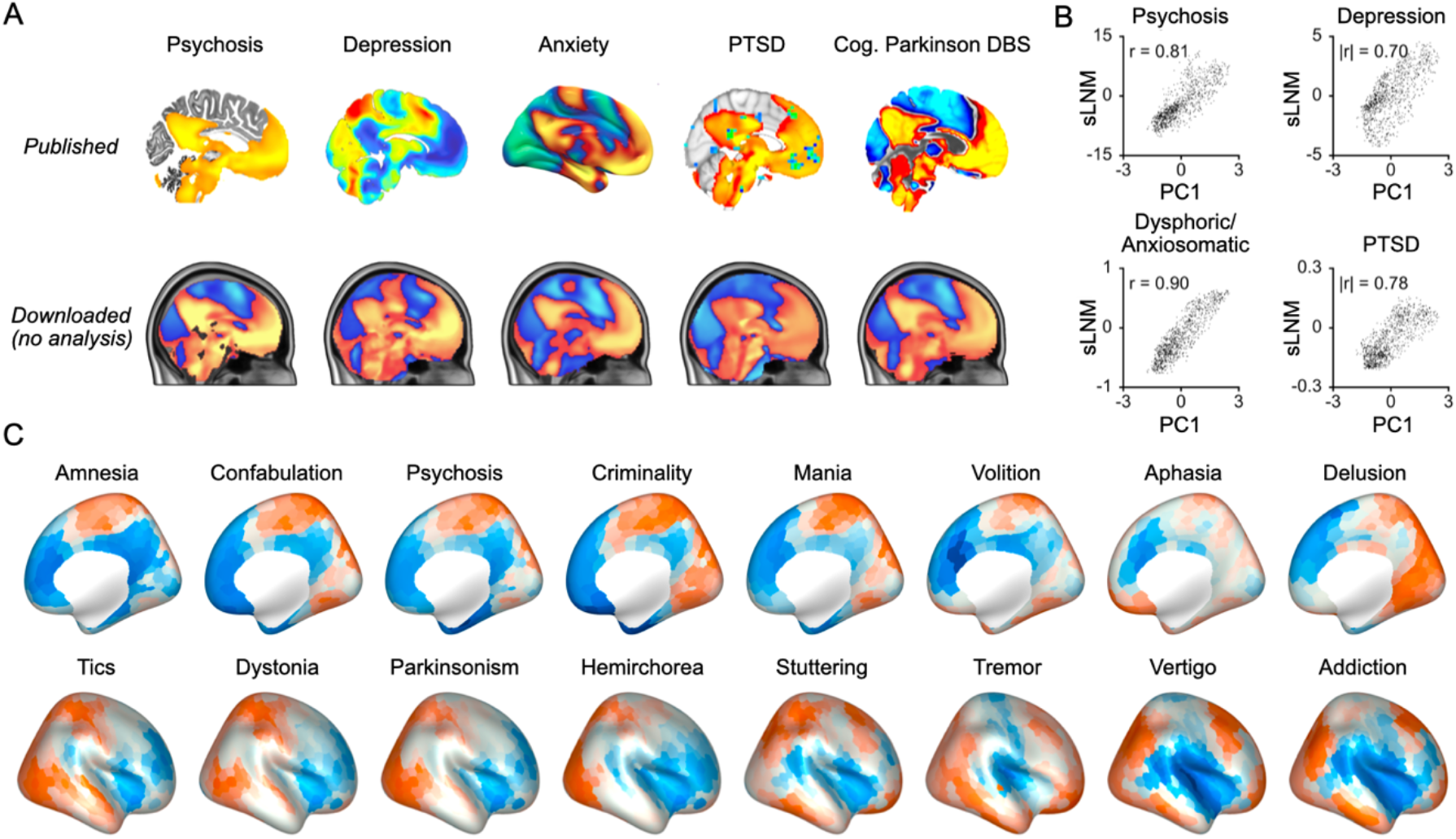
Specificity testing leaves the disease-specificity problem of LNM unresolved. **(A)** Lesion Network Mapping (LNM) specificity maps of five exemplary conditions and group-contrast maps as presented in the literature. The upper row (‘Published’) reproduces panels from independent publications [psychosis, depression, anxiety, PTSD, DBS induced cognitive decline in Parkinson’s disease^8-10, 15, 55^] showing LNM specificity maps resulting from specificity testing by comparing LNM lesion maps against those from control lesions, contrasting patient groups, and/or applying symptom LNM (sLNM). The lower row (‘Downloaded’) shows specificity maps downloaded directly from original publications [psychosis^8^; depression^17^; dysphoric-anxiosomatic^6^; PTSD^10^; DBS induced cognitive decline in Parkinson’s disease^9^; contrasts in the depression and PTSD maps are sign-inverted], illustrating highly similar LNM specificity maps across clinically unrelated conditions. Displayed maps were downloaded directly from the original LNM investigations and are shown without any further analysis. **(B)** Scatter plots of the relationship between specificity/sLNM maps and PC1 of the normative connectivity matrix C. These examples are also presented in the original study^3^. **(C)** Rows show examples of specificity LNM maps from 16 conditions discussed by Siddiqi et al.^5^ and others, derived using two-sample t-tests comparing lesions of interest with a large control lesion set. Upper and lower rows depict distinct classes of LNMs within the constrained set of recurrent specificity patterns defined by the connectivity matrix C.

These correlations are unusually high for neuroimaging studies. In fact, values of r = 0.6–0.8 approach a practical ceiling for LNM analyses, given that replacing the GSP1000 connectome with alternative datasets in a test-retest analysis, such as HCP^11^ or FC1000 data^12^, produces variations of comparable magnitude when the same lesion sets are analyzed (Supplementary Notes); this regularly leaves LNM maps for unrelated conditions statistically equivalent (Supplementary Notes). Indeed, correlations within this range are routinely presented in LNM studies as evidence of similarity and reproducibility of LNM networks, for example for glioma^13^, preclinical Alzheimer’s disease^14^, and others^8, 15-18^.

This overlap has been interpreted as reflecting a convergence on “degree”^5, 19, 20^, with statistical refinement by means of correcting for such “hub-related” or “connectome-driven bias” proposed as solutions to the raised concern. However, this interpretation has been cautioned against^3^, as such adjustments do not address the fundamental limitation. LNM linearly projects lesion locations *M* onto a single normative connectome *C*, constraining its outputs by the generic structure and low-dimensional patterns available in *C*^3, 4, 21^. This makes repetition of similar networks a default outcome of the method, with the matrix-average (“degree”) emerging as a recurrent motif alongside other canonical patterns when additional operations such as contrasts are applied^3^, often leaving similar networks to be attributed to unrelated conditions.

### Specificity testing

Although LNM sensitivity maps remain a primary reported result in LNM studies, including in recent studies (e.g.^13, 22-25^), *specificity testing* has been put forward as the key second step to establish anatomical specificity of the derived networks^5, 19, 20^, leading to the suggestion that the previous methodological investigation omitted this component^5, 15, 26-28^.

This incorrect impression may have arisen from the variable terminology used in the LNM literature. *Specificity testing* is often presented in LNM studies as a separate second stage of the analysis^5, 19^, while symptom-based LNM (*sLNM*) is described as a methodological extension or “next generation” of the method that further incorporates individual patient symptom scores or patient subgroups to derive symptom-specific brain networks^6, 10, 16-19^. This variable terminology (see also Supplementary Note S9^3^) may have made these procedures appear more distinct than they are: *specificity testing* and *sLNM* implement the same basic operation, contrasting lesion-derived connectivity profiles after projecting lesion information *M* through the same normative connectome *C*.

In LNM, the standard network, or sensitivity map, is obtained by selecting the functional connectivity (fc) profiles corresponding to lesions *M* from a normative connectome *C* and averaging them into a joint network^3-5^. Specificity testing applies the same operation to the lesion set of interest and to a control lesion set, and contrasts their connectivity profiles using a two-sample t-test^5^ or by permutation of the symptom/group labels^19^. In sLNM, lesion-derived fc maps are related to symptom scores or group labels using correlation or regression within a general linear model (GLM)^10, 18^. These are not independent analytical operations, but alternative formulations of the same group contrast: a two-sample t-test is a special case of the GLM, in which the design matrix encodes group membership and the group coefficient gives the difference between the two group means. sLNM therefore provides the more general formulation of specificity testing. Accordingly, performing specificity testing using a two-sample t-test between a set of lesions of interest and a set of control lesions, or applying sLNM to the combined lesion sets using an equivalent group-membership vector, produces identical results (r = 1.00).

The concern that specificity testing was not yet evaluated in the methodological investigation of LNM^5, 19^ therefore appears to reflect a difference in terminology rather than an omission of the underlying group-contrast operation. Relevant examples of specificity testing, group- and symptom-contrast testing were presented jointly within the broader sLNM framework (collectively marked by *“s”* in e.g. Figure 1, 4^3^ and Supplementary Notes of^3^, see also^4^). These and additional examples of reported specificity networks in recent LNM literature presented here (Figure 1A) clarify that LNM networks derived through the proposed step of specificity testing do not escape the raised limitation and continue to show repetition across unrelated conditions.

Below, we expand on the discussion of the methodological operations underlying sensitivity and specificity testing in LNM^3^, clarify their relationship to sLNM, and provide empirical examples showing that specificity testing does not provide anatomical specificity of derived disease and symptom circuits.

## Results

### Investigating the steps underlying sensitivity and specificity testing in LNM

As mentioned, the *sensitivity* step of LNM projects a set of lesions associated with a specific symptom or condition of interest (e.g. lesions associated with addiction^16^, psychosis^8^, cognitive impairment^19, 29^) onto a normative connectome, with their joint mean functional connectivity pattern interpreted as the circuit causally linked to the studied symptom^2, 30^. This is formalized by *LNM = ΣMC*, where *M* describes the normalized lesion matrix, and *C* describes a voxel- or region-wise normative functional connectivity matrix, resulting in fc maps similar to those derived by Lead-DBS (r = 0.94-0.98)^3^.

In *specificity testing*, the same operation is applied, but now for two lesion groups. The group of interest, *Group A*, samples one set of fc maps from the normative dataset (i.e. *M*_*A*_*C*), and a control set, *Group B*, samples another set of fc maps from the same normative matrix (i.e. *M*_*B*_*C*). The principal contrast of interest is the difference between the means of these two selected fc sets, *mean(M*_*A*_*C)* − *mean(M*_*B*_*C)* (similar to *Σ*, when *M* is normalized^3^). With both maps derived from the same connectivity matrix *C*, this step effectively involves comparing the means of two aggregations of rows from the same matrix *C*: *LNMspecificity = mean(rows of C matching Group A) − mean(rows of C matching Group B)*, that is, subtracting two sensitivity maps, *LNMspecificity = LNM*_*A*_ − *LNM*_*B*_. Figure 2 illustrates this procedure; interactive examples are presented at methods.lnmviewer.org. An inferential step can be added by standardizing this contrast by the within-group variability and evaluating it against a theoretical distribution, as in a two-sample t-test implementation of specificity testing^5^; alternatively, symptom- or group-label permutation can be used to generate an empirical distribution of random contrasts under reassigned labels. Both implementations yield similar specificity maps (r = 0.98).

**Figure 2.**
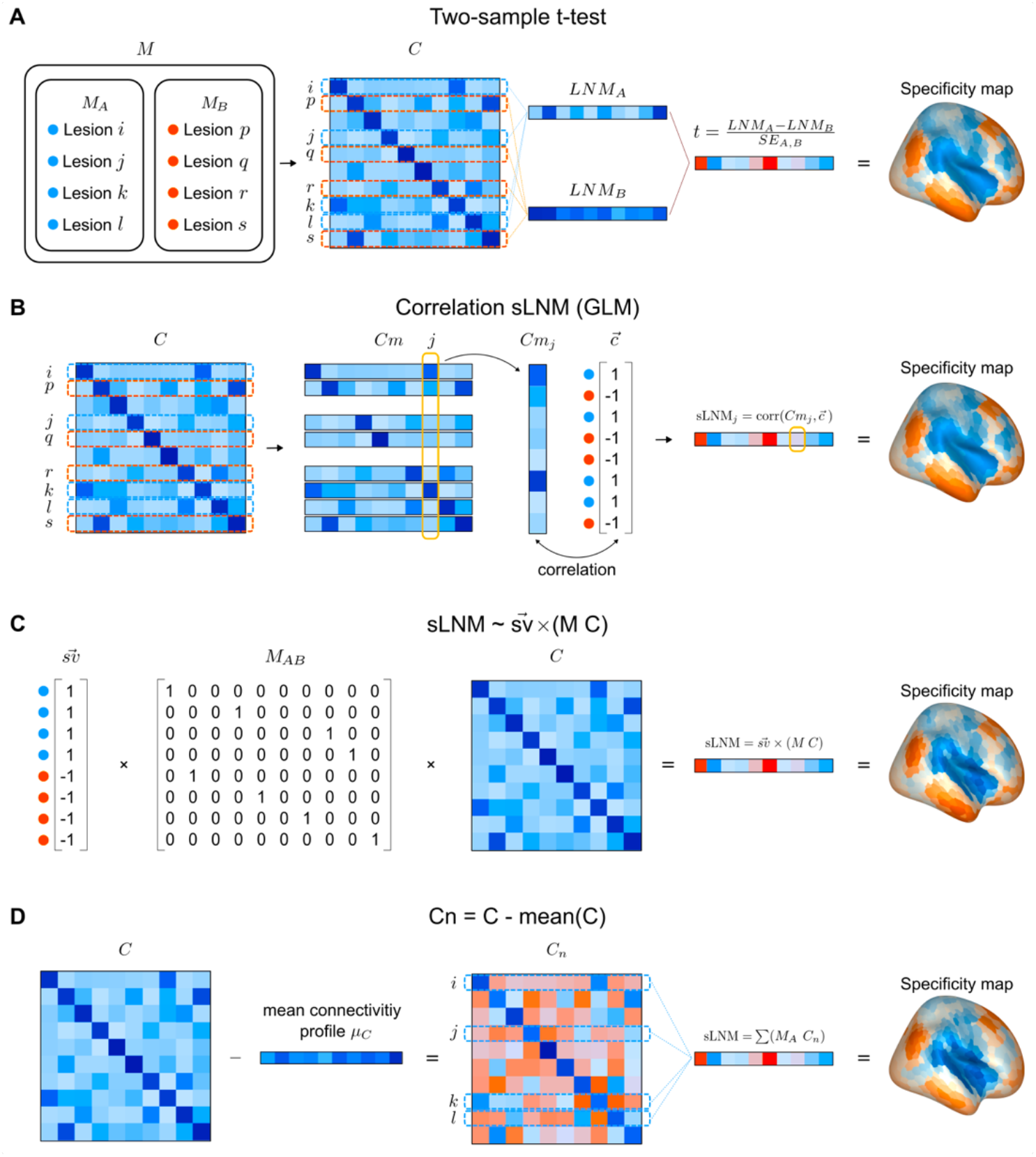
Methodological similarity of specificity testing, sLNM, and matrix replacement. Figure illustrates specificity testing using a two-sample t-test, its formulation within the symptom Lesion Network Mapping (sLNM) framework using a general linear model (GLM), and by means of replacing normative matrix C with a centered matrix Cn ∼ C − mean(C). **(A)** Specificity testing in LNM compares functional connectivity (fc) maps of the lesions M (left) of two groups, A and B. The lesion matrices M_A_ and M_B_ are projected onto C (middle), and the fc profiles corresponding to the two sets of rows of C are compared using a two-sample t-test (right). **(B)** The same group comparison can be formulated within the sLNM framework by defining a symptom vector sv as a binary contrast vector c labeling Group A as 1 and Group B as −1 and performing a regression analysis/GLM using c and M_AB_ C. **(C)** This group-contrast map can also be expressed using the approximation of sLNM ∼ svMC^3^. Normalizing the positive and negative entries of c for the number of lesions in M_A_ and M_B_, respectively, the contrast is obtained as cM_AB_ C. **(D)** In many LNM studies, Group B consists of a large and anatomically heterogeneous collection of control lesions. Under this setting, the LNM map of Group B approximates mean(C), or, after accounting for lesion size and anatomical lesion prevalence, mean(Cp), where Cp denotes a lesion-prevalence-weighted version of C. Because the operations applied to M and C are linear, the same specificity map can be derived by defining a matrix Cn = C − mean(C) by subtracting the mean connectivity profile from each row of C, and using this alternative matrix Cn in the standard LNM procedure, LNMspecificity ∼ ΣM_A_Cn. If desired, Cn can be scaled by the standard error of the contrast between Groups A and B to obtain values in the range of t-scores, or approximated from C more directly (Supplementary Notes).

### The control group in specificity testing

In many LNM studies, the control Group B comprises a large, heterogeneous set of control lesions, often drawn from broad lesion repositories spanning multiple conditions, such as the Harvard Lesion Repository^31, 32^. Because this control set samples a wide range of rows from *C*, its mean connectivity profile approaches the row-average profile of *C* itself^3^. Furthermore, when the nonuniform spatial distribution of real lesions is accounted for^5, 33, 34^, Group B can be even more closely approximated by the mean of a lesion-prevalence-weighted connectivity matrix (*Cp*). In this common setting, *mean(C)* closely resembles *mean(Group B)* (r = 0.77; r = 0.99 when using *Cp*), and *LNMspecificity ≈ mean(rows of C matching Group A) − mean(rows of C matching Group B)* becomes *LNMspecificity ≈ LNM*_*A*_ *− mean(C)*^3^.

### Reformulating specificity testing using an alternative matrix Cn

Because the above operations are weighted sums of connectivity profiles drawn from the same normative connectivity matrix *C*, the contrast can be equivalently obtained by first replacing matrix *C* with a centered matrix, *Cn* = *C − mean(C)*, where we subtract the row mean connectivity profile of *C* from each row of *C*, and then apply the standard LNM operation directly to *Cn* (for a further improved approximation, one can use *Cnp = C − mean(Cp)*). This effectively leaves specificity testing in LNM to *LNMspecificity ∼∑M*_*A*_*Cn*. Figure 2 presents a schematic representation. If desired, *Cn* can be further scaled to obtain values in the same range as values from the t-test (Supplementary Notes). Indeed, using *∑M*_*A*_*Cn* produces specificity maps highly similar to those obtained with the full proposed two-sample specificity t-test procedure^5^ on example lesion sets (*Cn:* r = 0.83 and even more precise, using *Cnp:* r = 0.96).

### The scope of specificity testing in LNM

This formulation, using a replacement matrix *Cn*, helps clarify the interpretation of the contrast examined by specificity testing in LNM. Two-sample t-tests and permutation-based models are commonly used in neuroimaging to assess differences between groups, for example between patients and controls. In LNM, the data for both Group A and Group B, and their contrast, are inherently generated from the same normative connectivity matrix *C*. Both groups inherit the dominant matrix-average pattern of *C* and specificity testing may attenuate a shared component (for example, “degree”); however, removing *mean(C)*, such as *Cn = C − mean(C)*, does not make the resulting contrast independent of *C*. Rather, matrix *Cn* remains entirely derived from *C* and retains much of its low-dimensional structure. Consequently, the contrast between Group A and Group B remains shaped by the principal axes of variation in *Cn*, and by those of the original connectome *C* (empirical examples are discussed in more detail below). Specificity networks, like sensitivity maps, remain therefore constrained by the standard patterns of *C*^3, 4, 21^, largely irrespective of the disease or symptom under investigation.

### The dimensionality of C constrains specificity maps

The consequences of this constraint can be demonstrated by examining specificity testing using an example connectome matrix *Cr*, whose variation is described by a single dominant axis, such as the first principal component of the connectome (PC1 represents a dominant motif explaining 39% of the variance in *C*). Using *Cr*, each row *i* can be written as *Cr*_*i*_ *= μ* + *a*_*i*_PC1, where *μ* is the mean connectivity profile of *C*, and *a*_*i*_ is the PC1 score of row *i*.

Performing a specificity test on *Cr* involves sampling one set of rows for Group A and another for Group B, with their fc maps contrasted by means of the two-sample t-test: *LNMspecificity = mean(rows of Cr matching Group A) − mean(rows of Cr matching Group B)*, which translates to *LNMspecificity* 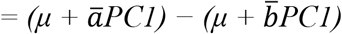, and further, *LNMspecificity* 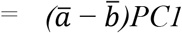, where ā and 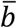 denote the mean PC1 scores of the lesions/rows selected by Group A and Group B, respectively.

When Group A and Group B sample identical or equivalent rows of *Cr*, 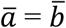, their contrast is naturally zero. When both groups sample randomly from *Cr*, the difference between ā and 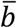 is expected to be small and to converge toward zero as sample size increases, with again no significant specificity contrast emerging. However, when a significant group difference is observed, it must necessarily arise from a difference in the groups’ scores along PC1, since *LNMspecificity = (*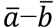*)PC1*. A resulting contrast may thus show significant results, but because all centered variation in *Cr* lies along a single axis, the resulting spatial contrast inevitably expresses a pattern that is proportional to *PC1*. Statistical significance in specificity testing may thus establish that the two lesion groups sample different parts of the connectome space, leading to significant voxels, but this does not establish a pattern or network that is disease-specific or independent of *C*; the resulting map can only express the pattern(s) present in the used matrix, i.e. PC1 when using *Cr* in the example.

The full matrix *C* naturally contains additional axes of variation (including *PC2, PC3, PC4*, etc.), and therefore permits a broader range of contrasts, but the same principle applies: specificity maps remain constrained by the effective dimensionality of *C* (∼3–4 dominant dimensions^3, 35-37^), and by the spatial axes those dimensions define. Because these components account for most of the variance in the normative connectome, specificity maps are largely restricted to the limited set of latent patterns defined by these gradients. Applying specificity testing to different lesion sets across multiple studies is therefore expected to produce maps that eventually converge on the same latent patterns. Similarly, thresholded specificity peaks in these maps are expected to recur in regions located at the extremes of these dominant gradients.

### Repetition of empirical LNM specificity maps across studies

We empirically examined specificity testing in LNM by reconstructing LNM specificity maps for 31 of the 34 conditions described by Siddiqi and colleagues^5^, covering 874 of the lesions tested^21^ (Supplementary Notes). For all lesions, fc maps were derived using the voxel-wise Lead-DBS and atlas implementation^3^, and compared with those from a large control lesion set (n = 1,000) using the recommended two-sample t-test procedure^5^, deriving specificity maps for 31 conditions.

Although specificity testing is proposed to ensure specificity to the symptom studied, overlap among the resulting specificity maps is substantial (|r| = 0.35, SD = 0.25): on average, the 31 tested conditions showed a network that overlapped at |r| > 0.6 with five other unrelated conditions (SD = 3.8; Figure 1C). Expanding this analysis with 40 additional lesion sets^3^ (Supplementary Notes) confirmed the observed overlap, with each of the 31 networks matching, on average, a pattern also observed in 11 other conditions (|r| > 0.6). This overlap is not unexpected, given that effectively the contrast examined is captured by *C − mean(C)*, as derived above. Much of the information in the specificity map can thus also be obtained by subtracting the mean connectivity profile *mean(C)* directly from the original sensitivity map, leaving the overlap structure across conditions largely unchanged (r = 0.86). The tendency of LNM maps to resemble one another^3^, argued to be an expected feature^5^, thus largely persists after specificity testing. While appearing distinct when presented individually, LNM specificity networks show substantial repetition and non-specificity when compared across studies.

The repetition of LNM specificity maps becomes further visible when we examine the shared underlying structure of specificity maps. A principal component analysis of 11 published specificity/sLNM maps downloaded directly from the original publications (without further analysis) shows that the first three principal components explain 85% of their joint variance (see also^38-40^ and Figure 1B for examples). This low-dimensional structure is also observed in the larger sets of 31 and 40 specificity maps (PC1–5 explain 90%). Notably, these patterns are naturally related to the main axes of variation of the used connectivity matrix *C* (PC1–3 of *C*, r = 0.69).

The recurrent nature of specificity results is also evident in the *specificity peaks* frequently reported in LNM studies. We revisited the literature overview of LNM studies performed in the original investigation^3^, and further collected MNI coordinates of 566 specificity peak locations reported for 78 conditions from 47 studies (Supplementary Methods). When compared across studies, documented peak locations show a systematic relationship with the standard patterns of *C* (PC1, p = 0.003, Supplementary Notes), with multiple studies reporting overlapping peak locations across unrelated conditions. For example, an inferior frontal specificity peak area reported for depression^17^ is also reported as a peak region by studies on post-stroke cognitive impairment^41^, out-of-body experience^42^, or sensorimotor behavior in stroke patients^43^, among other conditions (10–17 mm; Supplementary Notes, Table S2). This suggests that, while peak locations are often reported in individual LNM studies as key areas of a symptom-specific circuit and subsequently proposed as potential targets for neuromodulation, their recurrence across studies may primarily reflect standard patterns of *C* rather than disease-specific relationships.

## Discussion

Specificity testing in LNM statistically compares lesion maps between groups. Such testing provides an alternative benchmark against which the derived LNM network is tested and can readily yield statistically significant effects. However, it does not resolve the central methodological limitation of LNM: whether implemented through a two-sample t-test, sLNM, or related procedures, the resulting maps and their contrasts remain derived from the same normative connectivity matrix *C*. Contrasts between rows of *C* may attenuate the matrix-average pattern (i.e. the suggested “hub” or “degree” pattern^5^) and reweight its principal gradients, but they do not expand the underlying limited space of possible network outcomes.

The central issue is not a suggested omission of specificity testing from the methodological investigation of LNM, but a remaining dependence of LNM, sensitivity and specificity testing alike, on the used low-dimensional group connectivity matrix at the core of the method. This dependence makes recurrent network patterns across unrelated conditions a default outcome of the framework, rather than support for the existence of distinct symptom-specific circuits.

In the LNM literature, closely related variants of the framework are commonly presented under various labels, including *coordinate network mapping*^44^, *atrophy network mapping*^45^, *therapeutic network mapping*^46^, *tumor network mapping*^13^, and others^22, 47-49^, further combined with various names for the analytical steps performed, including sensitivity testing, specificity testing, sLNM, and others^13, 44, 45, 47, 50^. Although presented as distinct procedures, these approaches are mathematically related through overlapping operations applied to matrix *C*. Sensitivity-type operations, represented by *MC*, emphasize the matrix-average structure of *C*. Specificity-type operations, represented by centered or contrast-based forms of *MC*, may attenuate shared components, but instead emphasize remaining axes of variation of *C*^3, 4^. Their relationship is central to the methodological question: with both sensitivity and specificity maps arising from linear combinations of the same normative functional connectivity matrix *C*, their outcomes are overlapping and constrained by the recurring patterns of *C*.

In practice, this leaves current LNM applications often producing overlapping outcomes, with unrelated symptoms yielding non-intuitively similar circuits. For example, LNM applied to differentiate dysphoric and anxiosomatic symptoms in depression^6^ yields brain circuits highly similar to those reported for criminality^51^ (r = 0.81) or psychosis^8^ (r = 0.74, Figure 1B). Targeted case-control study designs involving carefully selected patient populations^19, 51, 52^ also do not escape this disease-specificity limitation. For example, analyses intended to ensure anatomically distinct circuits for post-stroke cognitive impairment^19^ by contrasting specific stroke patient subgroups yield similar ‘circuits’ to those obtained when testing arbitrary combinations of lesion sets, such as neglect syndrome versus freezing of gait (r = 0.82) or creativity versus dystonia (r = 0.86).

These examples emphasize an important distinction between statistical group differentiation and biological specificity in LNM. A specificity contrast can establish that two lesion groups significantly differ in how they sample the normative connectome structure; it does not, by itself, establish that the resulting spatial pattern constitutes a disease-specific circuit. When examined across studies, networks and peaks repeatedly localize to a limited set of recurring connectome-derived patterns, constraining the range of symptom- or disease-specific networks that can be identified. Alternative approaches to evaluating LNM maps, including null-model and regression frameworks, are proposed, and represent an active area of methodological development^53, 54^.

This discussion opens a constructive direction for future work. Rather than pursuing the goal of mapping each symptom or disorder to an isolated and unique ‘network’, future work may consider *connectome pleiotropy*: the principle that a small number of brain networks and gradients may shape a landscape of cognitive functions, clinical phenotypes, and symptoms.

From this perspective, overlap across conditions is not necessarily a nuisance to be eliminated. It may instead reflect a shared neural architecture of latent brain networks through which diverse clinical manifestations may emerge.

## Methods

### Lesion sets

Specificity maps were examined across 31 published psychiatric and neurological lesion network mapping (LNM) datasets, with 874 of the 1,090 reported lesions reconstructed by manually segmenting lesions from the original case reports and published figures and registering them to Montreal Neurological Institute (MNI) space using FSL (Supplementary Notes). Lesion sets were complemented with previously assembled LNM lesion datasets, resulting in a total of 2,177 lesion masks from 71 conditions. For studies without individual lesion masks, lesion prevalence maps or published MNI coordinates were reconstructed into standardized volumetric masks following the original LNM methodology. In addition, 11 published symptom-LNM (sLNM) maps were downloaded directly from the original publications for comparison. See Supplementary Notes for further details.

### LNM ceiling

LNM variability was assessed by recomputing LNM maps for identical lesion sets across using alternative normative functional connectivity datasets (GSP1000, HCP, FC1000, and 72 additional public connectome datasets; Supplementary Notes, for processing details, see^56^).

Across 71 lesion sets, mean spatial correlations between LNM maps for the same lesion set across alternative connectomes ranged from r = 0.66–0.89 (mean r = 0.81, SD = 0.05).

### Lesion Network Mapping

*Voxel-wise lesion network mapping* was performed using Lead-DBS v3.1^57^ with MATLAB R2024b and the required SPM12 dependencies. Analyses used the unmodified Lead-DBS implementation and the GSP1000 preprocessed normative functional connectome. Lesions were analyzed in MNI152 space using the lead_mapper interface and the cs_fmri_conseed_seed_tc.m function. Equivalently, atlas-based LNM involved mapping lesions to the 1,000 cortical regions of the Yeo-Schaefer1000 atlas^58^ and the Melbourne54 subcortical atlas^59^ and selecting the matching rows of the selected parcels from the group connectome matrix *C*. See Supplementary Notes for further details. *Specificity testing*. Voxel-wise and atlas-based specificity maps were computed by comparing lesion-derived functional connectivity maps from the lesion set of interest (Group A) with those from a heterogeneous control lesion set (Group B; *n* = 1,000) using two-sample t-tests. The same Group A − Group B contrasts were additionally implemented within a general linear model (GLM) using binary group labels, and compared with symptom Lesion Network Mapping (sLNM) implemented using binary symptom labels. Spatial similarity between specificity maps, GLM maps, and sLNM maps was assessed using Pearson correlation. Specificity maps were further compared with the mean connectivity profile and principal components of the normative connectome, as in^3^. See Supplementary Notes for further details.

