## SupplementaryNotes for "Assessing specificity testing in Lesion Network Mapping"

#### Data resources

*Lesions.* Access to the original lesion masks was requested but was denied due to data restrictions. We therefore reconstructed 874 of the 1,090 lesions discussed by Siddiqi et al.[1] by downloading the original case report publications for 31 of the 34 lesion sets discussed[2-26]. For three conditions, we were unable to locate the original lesions. Lesions were manually segmented from the published figures and/or original case reports, and lesion masks were registered to the Montreal Neurological Institute (MNI) brain template using the FMRIB Software Library (FSL)[27].

*Additional lesion sets.* The lesion dataset was extended with 28 additional lesion sets previously investigated using LNM and reported in the original work[28], for a total of 2,177 segmented lesions.

*Prevalence maps.* For eight conditions, original lesions were not published, and only lesion prevalence data were available. In these instances, the MNI slice(s) with prevalence values were sampled from the figures in the study, mapped to MNI space, and regional prevalence scores were extracted in the Melbourne54 and Yeo-Schaefer1000 atlas spaces. 100 lesion locations were sampled per condition according to the reconstructed prevalence distribution.

*Coordinate data.* For resources in which MNI coordinates of cortical and subcortical abnormalities were reported, coordinates were extracted from the main report where available. When the coordinate list was not provided, coordinates were extracted directly from the original studies where possible. Standardized volumetric masks were created by dilating the coordinate points by 5 voxels in all directions, following the original steps of the LNM papers[29]. In total, 710 brain coordinates were extracted for 10 conditions.

*Extended open lesion datasets.* The dataset was expanded with 1,495 publicly available lesion masks from stroke[30, 31], tumor[32], glioma[33, 34], and multiple sclerosis[35] databases.

Together, lesion masks served as a real-world distribution of lesions, from which heterogeneous control groups for comparison were drawn ($n$ = 1,000).

*Downloaded (s)LNM maps.* Eleven published specificity and symptom-LNM maps (sLNM) were downloaded directly from respective publications (see original work[28]), for depression[36], post-traumatic stress disorder[37], anxiety-depression symptoms[38], psychosis[26], impaired verbal memory in multiple sclerosis[39], depression-related symptoms in multiple sclerosis[40], political involvement[41], as well as DBS-derived or related networks for obsessive-compulsive disorder[42], cognitive decline in Parkinson's disease[43], tremor relief[18, 44], and Alzheimer's disease[45].

**LNM tools**

Voxel-wise LNM was performed using Lead-DBS v3.1[46] (source: https://www.lead-dbs.org/download-lead-dbs/) and MATLAB R2024b, along with the required SPM dependencies (source: https://www.fil.ion.ucl.ac.uk/spm/software/spm12/). No modifications to the source code of Lead-DBS were made. The installation instructions available in the Lead-DBS user guide (https://netstim.gitbook.io/leaddbs) were used (section: data description) to download and install the GSP1000 Preprocessed Connectome data[47] from the Harvard Dataverse (source: https://doi.org/10.7910/DVN/KKTJQC). LNM was conducted using the Lead-DBS lead_mapper GUI and the cs_fmri_conseed_seed_tc.m function. Lesions were processed using Lead-DBS in the MNI152 coordinate system.

**LNM variability**

Variability of LNM maps related to the normative dataset was assessed by recomputing maps for identical lesion sets (e.g., addiction, migraine, vertigo) while varying the normative functional connectivity matrix *C* across datasets, including GSP1000[46], a test-retest sample of the Human Connectome Project[48], the 1000 Functional Connectome Project[49], and 72 additional group connectomes from publicly available resting-state fMRI data resources[50], processed and mapped to the Yeo-Schaefer1000 and Melbourne54 atlases[51, 52]. LNM spatial variability was quantified by deriving LNM maps for all lesion sets using 75 alternative normative connectomes and assessing their similarity using spatial correlations, with mean similarity across lesion sets ranging from r = 0.66–0.89 (overall mean r = 0.81, SD = 0.05)[53].

**Equivalence testing of LNM networks**

*Equivalence testing of sensitivity networks.* Equivalence between LNM maps was assessed using a bootstrap-based root-mean-square deviation (RMSD) framework. Lesion-wise fc maps were generated for each lesion, and condition-specific LNM maps (Group A) were calculated by averaging across lesions. Lesion sets with >10 lesions were included, resulting in 62 lesion sets. Next, sensitivity LNMs were compared against all other conditions and differences between LNM maps were quantified using RMSD between group-average maps. Bootstrap resampling (1,000 iterations) incorporated lesion-level and connectome-level variability by resampling lesions from 75 alternative normative connectivity matrices (*Ck*). An equivalence margin (Δ) was defined as the 90th percentile of the within-group RMSD distribution. Maps were considered equivalent when the observed RMSD and, as a conservative criterion, the upper bound of its 90% bootstrap confidence interval remained below Δ, indicating that many LNM networks were statistically indistinguishable within the predefined equivalence margin. Under these conditions, on average, each LNM was found to be statistically equivalent to 12 (SD = 9) other conditions, with 57 of the 62 tested conditions showing statistical equivalence to at least one other condition (44 showing equivalence to 5 or more other conditions). For example, a network derived for lesions associated with addiction[15] was found to be statistically indistinguishable from the LNM networks proposed for epilepsy, vertigo, glioma, stuttering and 10 other conditions, indicating that no meaningful difference is left between these conditions.
*Equivalence testing of specificity networks.* Equivalence testing was further extended to the analysis of the LNM specificity maps. To this end, for each comparison, two lesion groups of interest (Group A1 and Group A2) were each contrasted against a shared reference lesion set Group B, consisting of 500 lesions randomly sampled from the full pool, excluding lesions that are part of Group A1 and Group A2. Specificity testing was performed, contrasting Group A1 to

Group B and Group A2 to Group B and the difference between the specificity map of Group A1 and Group A2 was again quantified as the root-mean-square deviation (RMSD) between the A1-versus-B and A2-versus-B specificity maps. Bootstrap resampling (1,000 iterations) incorporated lesion-level and connectome-level variability by resampling lesions within lesion sets A1 and A2 with replacement and randomly sampling from the set of alternative normative connectivity matrices (*Ck*). An equivalence reference distribution was generated by shuffling lesion labels between A1 and A2 while keeping group sizes fixed and retaining B as the reference. The equivalence margin (Δ) was defined as the 90th percentile of this distribution. Specificity maps were considered equivalent when the observed RMSD was below Δ. Under these conditions, on average, the specificity map of each condition showed equivalence with 28 (SD = 13) other conditions, with 61 of 62 showing statistical equivalence to at least one other condition and 57 of 62 conditions showing statistical equivalence to five or more other conditions.

**Sensitivity testing, specificity testing, group-contrast LNM, sLNM**

Across the literature, multiple labels are used for closely related linear operations underlying sensitivity and specificity testing in LNM. Standard LNM selects the functional connectivity profiles of lesion locations from a normative connectome $C$ and derives a consensus map either by averaging these profiles, $LNM = mean(MC)$, or by testing them voxel-wise against zero using a one-sample t-test. In specificity testing, this operation is extended to a contrast between two lesion groups. If $M_A$ and $M_B$ denote the lesion-selection matrices for GroupA and GroupB, the unscaled specificity map is $LNMspecificity = mean(M_AC) - mean(M_BC)$, or equivalently $LNMspecificity = (mean(M_A) - mean(M_B))C$. A voxel-wise two-sample t-test applies an additional normalization by the standard error of the group difference, $t(v) = [mean(M_AC(v)) - mean(M_BC(v))] / SE_{AB}(v)$, where $SE_{AB}(v)$ is the voxel-wise standard error of the GroupA versus GroupB contrast and $v$ indexes brain voxels. The same contrast can be expressed in a general linear model (GLM) framework as $Y = X\beta + \varepsilon$, where $Y = MC,$ and the group effect estimated through a contrast $c^T\beta$, with the contrast vector $c$ defining Group A and Group B comparison, such as $[0, 1]$ or $[1, -1]$ (differing only in scale). In *sLNM*, symptom labels or symptom scores *sv* are related to lesion connectivity profiles by correlation or regression. The operation can be written compactly as $sLNM \sim sv^TMC$, which, after centering *sv*, corresponds to the covariance between symptom scores and lesion connectivity at each voxel. We note that this formulation implicitly treats the standard deviation as constant across columns. For binary symptom labels, this numerator is algebraically equivalent, up to scaling and sign, to the Group A versus Group B contrast. Thus, standard sensitivity LNM, specificity testing, GLM-based group contrasts, and sLNM correlation analysis differ primarily in statistical framing and normalization, while their core operation involve averaging, weighting, or contrasting rows of the same normative connectome matrix $C$.

**Meta-analysis of specificity coordinates**

We revisited the performed literature review of LNM studies[28] and examined reported LNM specificity coordinates with respect to the dominant connectome axes. A total of 182 LNM empirical studies were reviewed, with MNI coordinates reported for specificity testing extracted from 47 studies for 78 reported conditions (566 peak clusters; Supplementary Table 1). Reported cluster size was incorporated where available (assuming standard 1 × 1 × 1 mm MNI voxels); a default cluster size of 50 voxels was assigned when this information was unavailable. When multiple coordinates were reported for a significant cluster, each cluster was represented by a single MNI coordinate, selected as the peak coordinate associated with the largest reported cluster extent. Extracted coordinates and reported cluster extent were used to create a binary specificity-overlap map, where voxels covered by one or more specificity clusters were counted once.

Two analyses assessed the relationship between specificity voxels and PC1. First, the voxel-wise PC1 map was divided into 20 percentile-based bins, and specificity density was calculated for each bin as the number of reported specificity voxels divided by the total number of brain voxels in that bin. Spearman correlation was used to test whether specificity density increased or decreased with PC1. Specificity density varied systematically along the gradient (Spearman rho = 0.60, $p = 0.006$), indicating structured alignment with the PC1 gradient axis. Second, permutation testing assessed whether specificity voxels were shifted toward positive or negative PC1 values. Across 10,000 iterations, the same number of voxels as in the binary specificity-overlap map was randomly sampled from the brain mask, and the mean signed PC1 value was calculated to generate a null distribution. The observed mean signed PC1 value across specificity voxels was compared with this distribution. Specificity voxels were significantly shifted toward positive PC1 values ($z = 34.7$, $p < 0.001$).

*Depression example.* Ten specificity peak coordinates reported for unrelated conditions nearest to a reported depression-specific peak in the right inferior frontal gyrus (MNI: [46, 4, 35]; for complete list, see Supplementary Table 2)[36] included peaks associated with processing of angry faces[54] (10.2 mm), post-stroke cognitive impairment[55] (two peaks: 10.4 and 14.5 mm), out-of-body experience[56] (11.6 mm), sensorimotor behavior in stroke patients[57] (two peaks: 12.7 and 16.4 mm), cervical dystonia[5] (two peaks: 13.0 and 16.9 mm), fundamentalism[58] (14.9 mm), and motor anosognosia[21] (16.9 mm). Similar patterns were observed for other depression-specific peak voxels; for example, Supplementary Table 3 shows another depression-specific peak in the left intraparietal sulcus (MNI: [−33, −53, 46]) with nearby specificity peaks reported across distinct clinical and behavioral domains.

### Tables

<table>
<tr><th>Condition name</th><th>Article</th><th>Peak specificity voxels</th><th>Reference</th></tr>
<tr><td>Peduncular hallucinosis</td><td>Network localization of neurological symptoms from focal brain lesions</td><td>(-48, -64, -14); (50, -66, -16); (-24, -40, 68); (-12, 32, 14); (-44, -48, -34); (14, -62, 40); (-18, -26, -6); (-38, 18, -8); (8, -34, -6)</td><td>Brain. 2015 Oct;138(Pt 10):3061-75. doi: 10.1093/brain/awv228</td></tr>
<tr><td>Coma</td><td>A human brain network derived from coma-causing brainstem lesions</td><td>(-40, 12, -16); (0, 38, 12); (-30, 8, -12); (-6, -62, 12); (-2, 38, 12); (-12, -16, -2); (18, -18, 2); (0, -14, 8); (-28, 2, -14); (28, 4, -12); (-6, 2, -8); (0, -24, -28); (-6, -36, -26); (8, -36, -32)</td><td>Neurology. 2016 Dec 6;87(23):2427-2434. doi: 10.1212/ WNL.0000000000003404</td></tr>
<tr><td>Cognition</td><td>Canceled connections: lesion-derived network mapping helps explain differences in performance on a complex decision-making task</td><td>(-56, 6, 44); (38, -2, 12); (-34, -36, 28); (-42, -44, -20); (38, -32, 28); (56, -56, -10); (56, 2, 50); (-40, -56, 2); (18, -56, -18); (-28, -42, -26); (44, -38, -18); (-36, -6, 8)</td><td>Cortex. 2016 May;78:31-43. doi: 10.1016/j.cortex.2016.02.002</td></tr>
<tr><td>Tremor relief</td><td>Identifying therapeutic targets from spontaneous beneficial brain lesions</td><td>(12, -18, -2); (-10, -18, 0); (12, -60, -20); (-34, -24, 0); (-10, -60, -18); (-26, -8, -2); (-32, -6, -2)</td><td>Ann Neurol. 2018 Jul;84(1):153-157. doi: 10.1002/ana.25285</td></tr>
<tr><td>Criminality</td><td>Lesion network localization of criminal behavior</td><td>(4, 66, 8); (54, 2, -38); (-30, 18, -18); (-46, -68, 40); (0, -52, 24); (24, -14, -20); (54, -60, 30); (-28, -14, -16); (-50, -68, 24); (-12, -76, -16); (-4, -56, 64); (-6, 8, 42); (-24, -2, 52); (26, 0, 56); (-30, -58, 48); (22, -72, 34); (30, -34, -50); (64, 10, 34); (-60, 10, 32); (-32, -36, -50); (38, -42, 40); (-26, -72, -56); (36, -72, -20)</td><td>Proc Natl Acad Sci U S A. 2018 Jan 16;115(3):601-606. doi: 10.1073/pnas.1706587115</td></tr>
<tr><td>Alien limb, Mutism</td><td>Lesion network localization of free will</td><td>Agency: (2, -50, 48) | Volition: (-20, 20, 34); (-10, -70, 30); (54, -48, -40); (-52, -46, -42)</td><td>Proc Natl Acad Sci U S A. 2018 Oct 16;115(42):10792-10797. doi: 10.1073/pnas.1814117115</td></tr>
<tr><td>Cognition</td><td>The neuronal network involved in self-attribution of an artificial hand: a lesion network-symptom-mapping study</td><td>(50, -38, 32); (32, 24, -10)</td><td>Neuroimage. 2018 Feb 1;166:317-324. doi: 10.1016/ j.neuroimage.2017.11.011</td></tr>
<tr><td>Dystonia</td><td>Network localization of cervical dystonia based on causal brain lesions</td><td>(48, -12, 30); (36, -32, 63); (-43, -22, 20); (7, 2, 63); (-55, -11, 12); (-35, -57, -17); (-34, -60, -11); (-36, -48, -17); (-37, -2, -5); (33, -23, 60); (20, -43, -5); (11, -1, 52); (66, 4, 14); (3, -11, 13); (2, 33, 20); (-25, 1, 10); (-27, 4, 0); (-26, -15, 0); (-4, 36, 14); (4, 42, 6); (22, -71, 10); (-18, -82, 42); (-23, -46, -5); (48, -6, 27)</td><td>Brain. 2019 Jun 1;142(6):1660-1674. doi: 10.1093/brain/ awz112</td></tr>
<tr><td>Alzheimer's disease, Frontotemporal dementia, Corticobasal syndrome, Aphasia</td><td>Network localization of heterogeneous neuroimaging findings</td><td>Alzheimer's disease: (36, -44, 16) | Delusions in Alzheimer's disease: (50, 2, 52); (62, -26, 52); (-62, 10, 10); (-64, -30, 46) | Corticobasal syndrome: (6, -40, 76); (-18, -12, 48) | Frontotemporal dementia: (32, -14, -10) | Progressive non-fluent aphasia: (-54, 16, 2)</td><td>Brain. 2019 Jan 1;142(1):70-79. doi: 10.1093/brain/awy292</td></tr>
<tr><td>Loss consciousness</td><td>Cortical lesions causing loss of consciousness are anticorrelated with the dorsal brainstem</td><td>(-36, -6, -12); (36, -2, 10); (-56, 14, -8); (10, -14, 38); (-20, 0, -18); (-42, -14, -34); (22, 0, -18); (-36, -42, 68); (-6, -10, 76); (-48, -10, 54); (2, -32, 2); (50, -38, -18); (34, -82, -14); (52, -18, -6); (46, -34, 64)</td><td>Hum Brain Mapp. 2020 Apr 15;41(6):1520-1531. doi: 10.1002/ hbm.24892</td></tr>
<tr><td>Migraine</td><td>Mapping migraine to a common brain network</td><td>(-22, -100, 12)</td><td>Brain. 2020 Feb 1;143(2):541-553. doi: 10.1093/brain/awz405</td></tr>
<tr><td>Alien limb, Corticobasal syndrome</td><td>Network localization of alien limb in patients with corticobasal syndrome</td><td>Corticobasal syndrome: (-22, -48, -60); (22, -50, -58); (-4, -14, 48); (36, -10, 62); (-26, -14, -66); (32, -42, 66) | Alien limb: (-14, -22, 40); (-12, -48, 56); (-26, -50, 68); (14, -24, 42); (12, -44, 56); (22, -50, 64); (24, 32, 52); (40, 16, 52); (18, 44, 44); (-50, -56, 48); (50, -62, 50); (0, -44, 36); (-2, -66, 38); (10, -58, 34)</td><td>Ann Neurol. 2020 Dec;88(6):1118-1131. doi: 10.1002/ ana.25901</td></tr>
<tr><td>Alzheimer's disease</td><td>Network localization of clinical, cognitive, and neuropsychiatric symptoms in Alzheimer's disease</td><td>Recall: (-12, -22, -8); (32, -6, -20); (-20, -40, 6); (28, -40, 6); (12, -58, -42); (-8, -26, 4) | Recognition: (-18, -34, -2); (34, -42, -2); (-36, -22, -20) | Delusion: (12, 20, 68); (38, 24, -14); (18, 10, 18); (-24, 52, 32); (-52, -62, -28); (-30, 52, -12); (-32, 22, -12); (66, -48, 38); (56, -54, -30); (42, -56, -28); (-14, 14, 68); (-2, 24, 18) | Alzheimer's disease: (46, -12, -24); (28, -22, 38); (-6, -2, 28); (44, -12, -24); (-6, 0, 28)</td><td>Brain. 2020 Apr 1;143(4):1249-1260. doi: 10.1093/brain/ awaa058</td></tr>
<tr><td>Depression</td><td>Brain stimulation and brain lesions converge on common causal circuits in neuropsychiatric disease</td><td>(-53, 41, 15); (48, 38, 23); (-46, 9, 31); (46, 4, 35); (-33, -53, 46); (34, -51, 46); (-57, -50, -8); (8, 24, -4); (-6, 58, 10)</td><td>Nat Hum Behav. 2021 Dec;5(12):1707-1716. doi: 10.1038/ s41562-021-01161-1</td></tr>
<tr><td>Cortical dysplasia</td><td>Categorizing cortical dysplasia lesions for surgical outcome using network functional connectivity</td><td>(0, 8, 32); (70, -26, 20); (-66, -32, 26); (-56, 8, -2); (-12, 68, -8); (-8, -50, 42); (-42, -82, 34); (40, -62, 26)</td><td>J Neurosurg Pediatr. 2021 Aug 27;28(5):600-608. doi: 10.3171/2021.5.PEDS20990</td></tr>
<tr><td>Autoscopic phenomena</td><td>Common and distinct brain networks of autoscopic phenomena</td><td>Autoscopy: (18, -62, 36); (-17, -62, 32); (-8, -68, -32); (-39, -50, -59); (-35, -75, -21); (8, -74, -32); (-59, -51, -27); (54, -62, -12); (56, -54, -26); (60, -47, -26); (51, -47, -15); (-48, -56, -5); (30, 0, 56); (-18, -15, -15); (2, 2, -2) | Heteroscopy: (-47, 29, 15); (-36, -5, -11); (-20, -15, -27) | Out of body: (60, -48, 33); (12, -56, 45); (14, -47, 41); (35, 6, 38); (33, 18, 26); (51, 23, 29); (39, 26, -14); (-63, -56, 26); (-17, -75, -32)</td><td>Neuroimage Clin. 2021;30:102612. doi: 10.1016/ j.nicl.2021.102612</td></tr>
<tr><td>Hallucinations</td><td>Lesions causing hallucinations localize to one common brain network</td><td>Hallucinations all: (1, -68, -25); (-19, -38, -48); (20, -39, -48); (63, -23, -2) | Hallucinations visual: (-24, 22, -5); (25, -23, -4) | Hallucinations auditory: (1, -50, -28)</td><td>Mol Psychiatry. 2021 Apr;26(4):1299-1309. doi: 10.1038/ s41380-019-0565-3</td></tr>
<tr><td>Obsessive-compulsive disorder</td><td>Potential optimization of focused ultrasound capsulotomy for obsessive compulsive disorder</td><td>(0, 32, 18); (-40, 36, 30)</td><td>Brain. 2021 Dec 16;144(11):3529-3540. doi: 10.1093/brain/ awab232</td></tr>
<tr><td>Parkinson's disease</td><td>A brain network for deep brain stimulation induced cognitive decline in Parkinson's disease</td><td>(28, 12, -20); (-40, -58, -36); (-4, -52, -58)</td><td>Brain. 2022 May 24;145(4):1410-1421. doi: 10.1093/brain/ awac012</td></tr>
<tr><td>Religiosity</td><td>A neural circuit for spirituality and religiosity derived from patients with brain lesions</td><td>(-2, -36, -10); (0, -44, -14); (-12, -38, -8); (14, -38, -10); (-2, -10, -12); (-24, -8, 0); (0, 32, 4); (-32, 10, -8); (32, -16, -6); (-6, -56, 14); (40, -56, -32); (-44, -88, -2); (42, -88, -8); (62, -38, 42); (36, 6, 14); (46, 54, 2); (46, -46, 60); (26, -36, 32); (-22, -74, -10); (60, -60, -6); (38, 62, 0); (24, -76, -10)</td><td>Biol Psychiatry. 2022 Feb 15;91(4):380-388. doi: 10.1016/ j.biopsych.2021.06.016</td></tr>
<tr><td>Cognition</td><td>Activation network mapping for integration of heterogeneous fMRI findings</td><td>Anger: (50, 16, -16); (-50, 14, -14); (-54, -2, 50); (50, -4, 40); (-4, 2, 64) | Disgust: (-18, 0, -42); (-46, -36, -20); (36, -46, -16); (46, -40, 14) | Emotion: (12, 8, -32); (-52, 16, -14); (36, 16, 24); (-4, 4, 64); (34, -4, 16); (26, -52, 52); (-16, -30, -2); (-2, 42, -32); (-10, -18, 44); (0, 6, 12) | Fear: (16, 10, -30); (-54, 0, 50); (0, 44, -34); (26, -52, 50) | Happy: (-52, 18, -16)</td><td>Nat Hum Behav. 2022 Oct;6(10):1417-1429. doi: 10.1038/ s41562-022-01371-1</td></tr>
<tr><td>Addiction</td><td>Brain lesions disrupting addiction map to a common human brain circuit</td><td>(38, 30, 6); (-34, 26, 8); (6, 12, 60); (68, -30, 34); (2, 6, -18); (64, -8, -28)</td><td>Nat Med. 2022 Jun;28(6):1249-1255. doi: 10.1038/ s41591-022-01834-y</td></tr>
<tr><td>Aphasia</td><td>Involvement of thalamocortical networks in patients with poststroke thalamic aphasia</td><td>Aphasia: (-22, 56, 4); (-42, 32, 42); (-52, -38, -18); (-44, -48, 34); (-10, -14, 0) | Dysarthria: (32, 52, 26); (14, 28, 22); (-34, -62, -28)</td><td>Neurology. 2023 Jan 31;100(5):e485-e496. doi: 10.1212/ WNL.0000000000201488</td></tr>
<tr><td>Apnea</td><td>Lesions causing central sleep apnea localize to one common brain network</td><td>(1, 15, 37); (29, -48, -50); (-34, -50, -48)</td><td>Front Neuroanat. 2022 Sep 29;16:819412. doi: 10.3389/ fnana.2022.819412</td></tr>
<tr><td>Stroke</td><td>Multimodal and multidomain lesion network mapping enhances prediction of sensorimotor behavior in stroke patients</td><td>(4, -24, -32); (10, 10, 66); (18, -16, 4); (-58, -28, 24); (-16, -14, 10); (-14, 14, 64); (-54, -60, 16); (-56, -60, 14); (-28, -18, 10); (42, -8, 34); (-6, -74, 38); (-28, -18, 6); (-52, -56, 16); (58, -6, 30); (54, -26, 26); (-54, -58, 4)</td><td>Sci Rep. 2022 Dec 27;12(1):22400. doi: 10.1038/ s41598-022-26945-x</td></tr>
<tr><td>Central poststroke pain</td><td>Network effects of brain lesions causing central poststroke pain</td><td>(-32, -92, -16); (26, -96, -12); (28, -54, 70); (6, -36, 74); (-30, -34, 52); (-30, -46, 52); (34, -20, 50); (-30, -22, 53); (-34, 16, -12); (40, 20, -12); (-3, 37, 20); (0, -18, 38); (-22, -94, -8); (22, -96, -5); (-30, -54, 60); (12, -24, 72); (-10, -40, 62); (32, -32, 58); (-10, -26, 74)</td><td>Ann Neurol. 2022 Nov;92(5):834-845. doi: 10.1002/ana.26468</td></tr>
<tr><td>Blindsight</td><td>Network localization of unconscious visual perception in blindsight</td><td>(-8, -32, 2); (10, -32, 4)</td><td>Ann Neurol. 2022 Feb;91(2):217-224. doi: 10.1002/ana.26292</td></tr>
<tr><td>Cognition</td><td>The brain mechanisms of self-identification & self-location in neurosurgical patients using virtual reality and lesion network mapping</td><td>Self-location: (-58, -48, 28); (20, -34, -10) | Self-identification: (42, 20, -14); (6, 24, 58)</td><td>medRxiv [Preprint]. 2022 Mar 03.22.22272566. doi: 10.1101/2022.03.22.22272566</td></tr>
<tr><td>Multiple sclerosis (memory)</td><td>Multiple sclerosis lesions that impair memory map to a connected memory circuit</td><td>(18, -32, 8); (-18, -34, 8); (-16, -62, 32); (20, -56, 28)</td><td>J Neurol. 2023 Nov;270(11):5211-5222. doi: 10.1007/ s00415-023-11907-8</td></tr>
<tr><td>Anosognosia</td><td>Network localization of awareness in visual and motor anosognosia</td><td>Cross-modal anosognosia: (-26, -59, 24); (30, -56, 21); (-37, -37, -9); (28, 35, -9) | Motor anosognosia: (12, 8, 72); (52, 19, -9); (-32, 41, 23); (26, 37, 21); (-48, 16, -7); (67, -36, 32); (-37, -54, -57); (22, 64, -55); (36, -50, -57); (-48, -51, -31); (36, 7, -12); (54, 3, 49); (-18, -58, -28) | Visual anosognosia: (-27, -59, 25); (6, 69, -8); (-27, -35, -14); (-6, 0, -16); (0, -44, -51); (-11, 27, -7)</td><td>Ann Neurol. 2023 Sep;94(3):434-441. doi: 10.1002/ana.26709</td></tr>
<tr><td>Disorientation</td><td>A neural circuit for spatial orientation derived from brain lesions</td><td>(2, -61, 10); (-15, -42, -50); (16, -41, -50); (61, 3, -17); (-63, -2, -13); (21, -6, -46); (-21, -7, -47); (12, 12, -29); (-49, -20, 61); (25, 16, -31); (-6, -3, 7); (55, -38, 42); (0, -44, -37)</td><td>Cereb Cortex. 2024 Jan 14;34(1):bhad486. doi: 10.1093/cercor/ bhad486</td></tr>
<tr><td>Fundamentalism</td><td>A neural network for religious fundamentalism derived from patients with brain lesions</td><td>(18, 50, -11); (46, 14, 46); (50, -58, 56); (4, 38, 40); (-35, -76, -45); (60, -28, -16); (-12, -86, -26); (-46, -76, -28); (-20, 52, -10); (2, -30, 30); (14, 6, 20); (2, -12, 29); (20, 23, 60); (26, 22, -11); (4, -8, 16); (-10, -36, 76); (22, -50, -56); (-54, -68, 12); (56, -66, 4); (-26, -42, -56); (-42, -40, -24); (30, -16, 62); (-12, -26, 0); (14, -19, 74)</td><td>Proc Natl Acad Sci U S A. 2024 Sep 3;121(36):e2322399121. doi: 10.1073/pnas.2322399121</td></tr>
<tr><td>Decision-making</td><td>Common neural dysfunction of economic decision-making across psychiatric conditions</td><td>(10, 8, 2); (12, 10, 0); (18, 8, -4); (14, -8, 6)</td><td>Neuroimage. 2024 Jul 1;294:120641. doi: 10.1016/ j.neuroimage.2024.120641</td></tr>
<tr><td>Amusia</td><td>Focal brain lesions causing acquired amusia map to a common brain network</td><td>(52, -42, 18); (40, -18, -2)</td><td>J Neurosci. 2024 Apr 10;44(15):e1922232024. doi: 10.1523/ JNEUROSCI.1922-23.2024</td></tr>
<tr><td>Anxiety</td><td>Heterogeneous brain atrophy sites in anxiety disorders map to a common brain network</td><td>(-21, 3, -21); (24, 6, -21)</td><td>Depress Anxiety. 2024 Apr 2;2024:3827870. doi: 10.1155/2024/3827870</td></tr>
<tr><td>Alice in Wonderland Syndrome</td><td>Lesions causing Alice in Wonderland Syndrome map to a common brain network linking body and size perception</td><td>(42, 71, 16); (-57, -51, 37)</td><td>Ann Neurol. 2024 Oct;96(4):662-674. doi: 10.1002/ana.27015</td></tr>
<tr><td>Ataxia</td><td>Localization and network connectivity of lesions causing limb ataxia in patients with stroke</td><td>(10, -64, -38); (-6, -64, -36)</td><td>Neurology. 2024 Sep 24;103(6):e209803. doi: 10.1212/ WNL.0000000000209803</td></tr>
<tr><td>Neurogenic stuttering</td><td>Localization of stuttering based on causal brain lesions</td><td>Neurogenic stuttering (literature): (-29, 0, -7); (30, -5, -10); (-38, 21, 11); (27, -75, 43); (-23, -75, 47); (57, -59, -16); (35, -49, 67) | Neurogenic stuttering (clinical): (-31, -7, -6); (31, 0, -15); (-6, -19, -36); (39, -22, 2); (-26, -1, -36); (-11, -21, -1); (-14, -58, -24)</td><td>Brain. 2024 Jun 3;147(6):2203-2213. doi: 10.1093/brain/ awae059</td></tr>
<tr><td>Gait</td><td>A human gait circuit derived from brain lesions and deep brain stimulation</td><td>Gait DBS: (-34, -18, -34); (-28, -14, -32); (26, -12, -34); (44, -16, -38); (-22, 12, -10); (16, 24, -18); (-16, 24, -14); (-8, -30, -20); (0, -46, -34); (26, -84, -18); (-20, -70, 64); (24, -94, -12); (32, -92, -6); (-16, -94, -14); (-28, -92, -8); (24, -58, 72); (28, -84, 46); (-12, -56, 76) | Gait lesion: (-20, -22, 8); (20, -20, 12); (20, -4, -38); (6, -16, -8); (-4, -16, -8); (-26, -14, -36); (2, 36, 4); (-14, -50, -38); (16, -56, -36); (4, -28, -12); (-54, -68, 2); (-50, -76, 20); (58, -64, -4) | Gait combined: (-24, -12, -34); (6, -10, -8); (52, -8, -42); (2, 26, 10); (0, -42, -12); (14, 4, 0); (16, -50, -34); (-8, -16, 18); (-12, -2, 0); (10, -14, 18); (0, -26, -40); (-4, -32, 22); (-24, -66, 66); (46, -62, -12); (-46, -48, -16)</td><td>medRxiv [Preprint]. 2025 Mar 30:2025-03. doi: 10.1101/2025.03.28.25324852</td></tr>
<tr><td>Depression treatment</td><td>Distinct antidepressant therapies act on a common brain network</td><td>(60, 6, -12); (-54, 0, -12); (-27, -3, -21)</td><td>Nat Commun. 2026 Jan 3;17(1):1176. doi: 10.1038/ s41467-025-67945-5</td></tr>
<tr><td>Political involvement</td><td>Effects of focal brain damage on political behaviour across different political ideologies</td><td>Conservative: (-26, 22, 40); (30, -51, 5); (24, 15, 36); (35, -30, 1); (-34, -79, 52) | Democrat: (-36, -65, 55); (48, -27, 2) | Liberal: (60, -48, 10); (39, 0, -16); (-52, -33, 0); (0, -76, 55); (35, 4, -40); (-49, 10, -17) | Political: (-28, 14, 56); (-38, -70, 47); (-5, -70, 49); (-7, -39, 38); (-33, 40, 44); (-10, -22, 16); (55, -30, 4); (-47, -30, 3); (26, -3, -13); (-28, -46, 8); (41, 31, -2); (-33, -9, 26); (-63, 16, -16) | Republican: (-38, 26, 52); (-6, -37, 36); (-29, -56, 8); (-32, 39, -8)</td><td>Brain. 2025 Mar 21:awaf101. doi: 10.1093/brain/awaf101</td></tr>
<tr><td>Schizophrenia</td><td>Heterogeneous patterns of brain atrophy in schizophrenia localize to a common brain network</td><td>Schizophrenia: (-48, -16, 10); (56, -2, 8); (-4, -12, 42); (8, -62, -34); (4, 24, 24) | Schizophrenia vs others: (-28, 6, -14); (-22, -4, -12); (-28, 6, -14); (30, -50, 30); (46, 0, -34); (-26, 38, 18); (10, 70, 32); (-30, 8, -18)</td><td>Nat Ment Health. 2025 Jan;3(1):19-30. doi: 10.1038/ s44220-024-00348-5</td></tr>
<tr><td>Functional and somatic symptoms</td><td>Lesion network localization of functional and somatic symptoms</td><td>Functional: (14, 8, -8); (-12, 4, -8); (8, -14, -18); (20, 4, 26) | Somatic: (24, 44, -20) | Functional and somatic: (6, 0, 6); (24, 38, -22); (-26, 38, -24)</td><td>medRxiv [Preprint]. 2025 Mar 7:2025.03.06.25323494. doi: 10.1101/2025.03.06.25323494</td></tr>
<tr><td>Cognitive impairment</td><td>Post stroke cognitive impairment: more than a lesion-symptom model</td><td>(24, -80, 6); (32, -6, 26); (20, -6, -10); (56, 4, 32); (66, -36, 30); (60, 16, 6); (32, -2, 14); (30, -18, -16); (26, -4, 10); (-30, 6, 12); (-30, -20, 26); (48, -10, 0); (22, -18, 18); (44, -10, 32); (44, -16, 32); (-42, -20, 8); (-44, -10, -6); (-34, 2, 22); (-58, 30, 10); (-36, 6, -4); (-36, -22, -12); (-58, 20, 24); (-40, -40, 2); (-40, -36, -6)</td><td>medRxiv [Preprint]. 2025.02.19.25322521; doi: 10.1101/2025.0 2.19.25322521</td></tr>
<tr><td>Tuberous sclerosis complex-related epilepsy</td><td>Prognostic application of lesion network mapping to epilepsy surgery outcomes in pediatric tuberous sclerosis complex</td><td>(-34, -36, -28); (41, -40, -19); (-40, -18, -6); (28, 2, 2); (46, 6, -2); (-38, 6, 6); (20, -64, 18); (40, -76, 30); (6, -38, 78); (-10, -12, 6); (-10, 44, 20); (-8, -62, 22); (-36, 16, 28); (-10, -96, 8); (-38, -6, 44); (-4, 22, 50); (2, 20, 52)</td><td>Epilepsia. 2025 Mar 5. doi: 10.1111/epi.18320</td></tr>
<tr><td>Aphasia</td><td>Thalamic disconnection from prefrontal cognitive control networks contributes to thalamic aphasia</td><td>(44, -52, -42); (32, -58, -46); (-52, 12, -30); (-44, -2, -44); (6, -52, -40); (48, 12, -38); (-22, -76, -28); (44, -80, -30); (-50, 26, 4); (-4, 34, -10); (32, 22, -16); (50, 30, -8); (-12, 24, 60); (46, -38, -2); (-14, 12, 12); (16, 14, 8); (10, 58, 6); (-48, -62, 42); (58, -60, 32); (-8, -50, 34); (-2, -18, 38); (38, 24, 40)</td><td>Brain Commun. 2025 May 16;7(3):fcaf191. doi: 10.1093/ braincomms/fcaf191</td></tr>
<tr><td>Aphantasia</td><td>Visual mental Imagery and aphantasia lesions map onto a convergent brain network</td><td>(-38, -56, -18)</td><td>Cortex. 2026 Apr;197:1-12. doi: 10.1016/j.cortex.2026.01.009.</td></tr>
</table>

***Supplementary Table 1. Meta-analysis of reported peak specificity coordinates from LNM studies****. Peak specificity coordinates across 78 conditions (first column) were extracted from 47 LNM studies identified from a review of 182 published LNM studies. For each study (second column: article title; fourth column: reference), the reported peak coordinates associated with condition-specific findings are listed in MNI space (x, y, z; third column). When multiple conditions were reported within a single study, peak coordinates are annotated by condition to distinguish their respective associations.*

| Rank | Paper | Condition | x | y | z | Distance (mm) |
|---|---|---|---|---|---|---|
| 1 | PENG_2022 | Anger | 50.0 | −4.0 | 40.0 | 10.25 |
| 2 | HUYGELIER_2025 | Cognitive impairment | 56.0 | 4.0 | 32.0 | 10.44 |
| 3 | BLONDIAUX_2021 | Out of body | 35.0 | 6.0 | 38.0 | 11.58 |
| 4 | JIMENEZ_2022 | Sensorimotor | 42.0 | −8.0 | 34.0 | 12.69 |
| 5 | CORP_2019 | Dystonia | 48.0 | −6.0 | 27.0 | 12.96 |
| 6 | HUYGELIER_2025 | Cognitive impairment | 44.0 | −10.0 | 32.0 | 14.46 |
| 7 | FERGUSON_2024 | Fundamentalism | 46.0 | 14.0 | 46.0 | 14.87 |
| 8 | JIMENEZ_2022 | Sensorimotor | 58.0 | −6.0 | 30.0 | 16.4 |
| 9 | CORP_2019 | Dystonia | 48.0 | −12.0 | 30.0 | 16.88 |
| 10 | KLETENIK_2023b | Motor anosognosia | 54.0 | 3.5 | 49.9 | 16.92 |

***Supplementary Table 2. Specificity peaks closest to the depression-specific right inferior frontal gyrus peak.*** *The table lists the ten specificity peak coordinates nearest to the depression-specific peak reported in the right inferior frontal gyrus by Siddiqi et al.[36] (MNI: [46, 4, 35]). The table shows the rank of each peak by proximity to the depression peak (first column), source study (second column), associated condition (third column), MNI x, y, and z coordinates (fourth to sixth columns), and Euclidean distance from the depression-specific peak in millimeters (seventh column). Distances ranged from 10 to 17 mm.*

| Rank | Paper | Condition | x | y | z | Distance (mm) |
|---|---|---|---|---|---|---|
| 1 | DARBY_2018 | Criminality | −30.0 | −58.0 | 48.0 | 6.16 |
| 2 | KIM_2022 | Pain | −30.0 | −46.0 | 52.0 | 9.7 |
| 3 | TETREAULT_2020 | Corticobasal syndrome vs. Progressive supranuclear palsy | −22.0 | −54.0 | 54.0 | 13.64 |
| 4 | KIM_2022 | Pain | −30.0 | −54.0 | 60.0 | 14.35 |
| 5 | SIDDIQI_2025 | Democrat | −36.0 | −65.0 | 55.0 | 15.3 |
| 6 | STOCKERT_2022 | Aphasia | −44.0 | −48.0 | 34.0 | 17.03 |

| | | | | | | |
|---|---|---|---|---|---|---|
| 7 | TETREAULT_2020 | Alien limb | −50.0 | −56.0 | 48.0 | 17.38 |
| 8 | SIDDIQI_2025 | Political intensity | −38.0 | −70.0 | 47.0 | 17.75 |
| 9 | STOCKERT_2025 | Aphasia | −48.0 | −62.0 | 42.0 | 17.94 |
| 10 | KIM_2022 | Central post-stroke pain | −30.0 | −34.0 | 52.0 | 20.15 |

***Supplementary Table 3. Specificity peaks closest to the depression-specific left intraparietal sulcus peak.*** *The table lists the ten specificity peak coordinates nearest to the depression-specific peak reported in the left intraparietal sulcus by Siddiqi et al.[36] (MNI: [−33, −53, 46]). The table shows the rank of each peak by proximity to the depression peak (first column), source study (second column), associated condition (third column), MNI x, y, and z coordinates (fourth to sixth columns), and Euclidean distance from the depression-specific peak in millimeters (seventh column), ranging from 6 to 20 mm.*
